# Switching between external and internal attention in hippocampal networks

**DOI:** 10.1101/2022.12.20.521285

**Authors:** Craig Poskanzer, Mariam Aly

**Author notes:** Please address correspondence to: Craig Poskanzer.

## Abstract

Everyday experience requires processing external signals from the world around us and internal information retrieved from memory. To do both, the brain must fluctuate between states that are optimized for external vs. internal attention. Here, we focus on the hippocampus as a region that may serve at the interface between these forms of attention, and ask how it switches between prioritizing sensory signals from the external world vs. internal signals related to memories and thoughts. Pharmacological, computational, and animal studies have identified input from the cholinergic basal forebrain as important for biasing the hippocampus towards processing external information, whereas complementary research suggests the dorsal attention network (DAN) may aid in allocating attentional resources towards accessing internal information. We therefore tested the hypothesis that the basal forebrain and DAN drive the hippocampus towards external and internal attention, respectively. We used data from 29 human participants (17 female) who completed 2 attention tasks during fMRI. One task (“memory-guided”) required proportionally more internal attention, and proportionally less external attention, than the other (“explicitly instructed”). We discovered that background functional connectivity between the basal forebrain and hippocampus was stronger during the explicitly instructed vs. memory-guided task. In contrast, DAN-hippocampus background connectivity was stronger during the memory-guided vs. explicitly instructed task. Finally, the strength of DAN-hippocampus background connectivity was correlated with performance on the memory-guided but not explicitly instructed task. Together, these results provide evidence that the basal forebrain and DAN may modulate the hippocampus to switch between external and internal attention.

**Significance Statement:** How does the brain balance the need to pay attention to internal thoughts and external sensations? We focused on the human hippocampus, a region that may serve at the interface between internal and external attention, and asked how its functional connectivity varies based on attentional states. The hippocampus was more strongly coupled with the cholinergic basal forebrain when attentional states were guided by the external world rather than retrieved memories. This pattern flipped for functional connectivity between the hippocampus and dorsal attention network, which was higher for attention tasks that were guided by memory rather than external cues. Together, these findings show that distinct networks in the brain may modulate the hippocampus to switch between external and internal attention.

## Introduction

While navigating daily life, we must balance paying attention to sensory stimuli streaming in from the external world and paying attention to our internal thoughts and memories. To accomplish this, the brain must dynamically shift between states in which attention is externally vs. internally oriented (Chun et al., 2011; Verschooren et al., 2019a, 2019b; Li et al., 2022). This flexible reorienting of attention allows us to acquire new information and leverage previously stored information, which is critical for interacting with our environment, making decisions, and planning future actions (Lepsien & Nobre, 2006).

What neural mechanisms allow the brain to efficiently switch between external and internal processing? Although a burgeoning line of research addresses this question (Honey et al., 2017; Verschooren et al., 2019a, 2019b; Li et al., 2022; Isenburg et al., 2023), much of this work focuses on attentional networks. Here, we test how attentional networks might differentially interact with the hippocampus to switch between internal and external states.

Although the majority of literature on the hippocampus explores its role in episodic and relational memory, recent work has demonstrated its involvement in external attention. For example, hippocampal activity patterns represent top-down attentional goals (Aly & Turk-Browne, 2016a, 2016b; Córdova et al., 2019). Moreover, individuals with medial temporal lobe damage are impaired on relational attention tasks that require spatial processing (Ruiz et al., 2020). The hippocampus is also involved in guiding external attention by leveraging information from memory (Hutchinson & Turk-Browne, 2012; Günseli & Aly, 2020). Thus, the hippocampus is ideally situated to coordinate internal and external attention. Indeed, computational and rodent work suggests that the hippocampus oscillates between attentional states that prioritize processing either external or internal input (Honey et al., 2017; Tarder-Stoll, Jayakumar, et al., 2020; Kerrén et al., 2018). The peak vs. trough of local theta cycles in the hippocampus may increase the strength of external and internal input respectively (Hasselmo et al., 2002; Honey et al., 2017). To better understand these state switches, it is necessary to explore what drives this switching, and which brain areas bias the hippocampus towards external vs. internal attention.

One area that may bias hippocampal switching is the basal forebrain, which provides acetylcholine to the hippocampus (Newman et al., 2012). The basal forebrain promotes external attention through cholinergic projections across the cortex (Villano et al., 2017). Within the hippocampus, acetylcholine strengthens afferent input about the external world and suppresses recurrent connections that support internally oriented processing (Hasselmo 2006; Tarder-Stoll, Jayakumar, et al., 2020; Decker & Duncan, 2020). At a behavioral level, acetylcholine supports encoding (Newman et al., 2012), and cholinergic agonists benefit performance on a hippocampally mediated external attention task (Ruiz, Thieu, & Aly, 2021). This combination of animal, computational, and human behavioral work provides compelling evidence that the basal forebrain may bias the hippocampus into an externally oriented state.

Conversely, the dorsal attention network (DAN) may bias the hippocampus toward an internally oriented state. The DAN is recruited when accessing long-term memories to direct attention (Stokes et al., 2012) and dorsal parietal damage (overlapping with the DAN) impairs episodic recall (Berryhill et al., 2007). More generally, dorsal parietal regions, overlapping with the DAN, may allocate attentional resources towards mnemonic processing (Cabeza, 2008; Cabeza et al., 2008; Wagner et al., 2005). This “attention to memory” (AtoM) hypothesis suggests that the DAN may interact with the hippocampus to access information from memory, and may thus be an important network for biasing the hippocampus towards internal attention.

Here, we evaluate this two-fold hypothesis in which the basal forebrain and DAN dynamically interact with the hippocampus to bias external and internal attentional states respectively. We leverage two attention tasks that vary in their proportional demands on external vs. internal attention. We hypothesize that hippocampal functional connectivity with the basal forebrain and DAN will switch based on attentional goals, with higher DAN-hippocampus connectivity and lower basal forebrain-hippocampus connectivity for tasks that require proportionally more internal, and proportionally less external, attention.

Testing this hypothesis can be challenging because many external attention tasks do not recruit the hippocampus (Aly & Turk-Browne, 2017; Dudukovic et al., 2011; Yamaguchi et al., 2004). Furthermore, a comparison of tasks that vary in internally vs. externally oriented demands would ideally keep stimuli and motor demands identical while varying only participants’ attentional states, adding further constraints on the types of tasks that can be used. To address these challenges, we used a dataset collected during attention tasks that reliably recruit the hippocampus, with hippocampal engagement replicated across multiple studies (Aly & Turk-Browne, 2016a; Aly & Turk-Browne, 2016b; Günseli & Aly, 2020; Ruiz et al., 2020). The key feature of these tasks is that they require attention to *relational* information, and particularly the ability to identify higher-order similarities between images that are not perceptually identical (see Methods). Although the tasks are complex, stimuli and motor demands are identical across conditions, and their reliable recruitment of the hippocampus was important for addressing our question.

## Methods

To test these hypotheses, we analyzed a dataset (Günseli & Aly, 2020) in which participants performed 2 attention tasks, one in which they searched for rooms with the same spatial layout and one in which they searched for paintings with a similar artistic style. These 2 attentional states each took place in 2 conditions: memory-guided attention and explicitly instructed attention. These conditions were identical in terms of stimuli and motor demands; they differed only in whether participants selected their attentional state (art or room attention) based on their memory for images in the preceding trial (“memory-guided” condition) or were assigned an attentional state at the beginning of each trial (“explicitly instructed” condition). These attentional states and task conditions strongly modulate the hippocampus (Günseli & Aly, 2020; also see Aly & Turk-Browne, 2016a, 2016b; Ruiz et al., 2020) but differ in their proportional demands on internal vs. external attention. This allowed us to examine how hippocampal network connectivity patterns vary when one task requires more internally oriented attention, and less externally oriented attention, than another.

### Participants

Participants, stimuli, and tasks were previously described in Günseli & Aly (2020) and are summarized again here. A total of 30 participants were recruited from the Columbia University community. Participants provided written, informed consent and were paid $72 for their participation. As in Günseli & Aly (2020), we analyzed data from 29 participants, excluding 1 whose performance was three standard deviations below average on the memory-guided attention task. The included participants (17 female; 1 left-handed; all normal/corrected-to-normal vision) were 18-35 years old (M = 26; SD = 4.07) and had 13-21 years of education (M = 17.1; SD = 2.2). Study procedures were approved by the Columbia University Institutional Review Board.

### Stimuli

A total of 141 images were created for the main fMRI task by combining a picture of a 3D-rendered room with an image of a painting. An initial 20 virtual rooms were rendered using Sweet Home 3D (sweethome3d.com). They were designed to have a unique combination of shape, furniture, and wall color. After capturing an image of each room, the viewpoint was rotated 30° (50% of rooms were rotated clockwise and 50% counterclockwise). Next, these rotated images were edited while preserving their spatial layout: the wall color was changed and all furniture was substituted with a different piece of the same category in the same position (e.g. substituting a couch with a different couch). A second image was then captured of this spatially consistent room, which we refer to as a “room match” to the original image.

20 painting images were initially selected from the Google Arts & Culture Project (https://artsandculture.google.com/). Each art image was then paired with an “art match” by selecting a second painting by the same artist. These 2 paintings were thus stylistically similar to each other, while containing unique features. To create a combined art/room image, a painting image was superimposed on a room image to appear as if it were hanging on one of the visible walls.

A set of images were then selected to be “stay” or “switch” cues. The presence of a “stay” cue in a given trial in the memory-guided condition would indicate that the participant should stay in the same attentional state (art or room) on the following trial; the presence of a “switch” cue would indicate that they should switch to the other attentional state on the following trial (**Figure 1A**). Stay/switch cues were created by selecting 2 rooms (1 stay cue and 1 switch cue) and 2 paintings (1 stay cue and 1 switch cue). Each room cue was paired with 3 selected paintings to form 3 room “stay” images and 3 room “switch” images; similarly, each painting cue was paired with 3 room images to form 3 painting “stay” images and 3 painting “switch” images. To create parity in the non-attended “background” of the stay/switch cues (i.e., “background” paintings for room stay/switch cues and “background” rooms for painting stay/swich cues), the same 3 paintings were used for both room “stay” and “switch” cues, and the same 3 rooms were used for both art “stay” and “switch” cues. This ensured that participants could not identify whether a given (art or room) cue was “stay” or “switch” based on the unattended “background” feature (room or art, respectively). The non-attended “backgrounds” of the stay/switch cues were also used to make art/room images that were not “stay” or “switch” cues. In particular, each of the 3 room backgrounds for art “stay” and “switch” cues was also paired with 2 painting images that were not “stay” or “switch” cues; and each of the paintings paired with room “stay” and “switch” cues was also paired with 2 rooms that were not “stay” or “switch” cues. This ensured that participants could not use these “background” features to identify that a “stay” or “switch” cue had been presented. In sum, this procedure resulted in 12 images in which stay/switch cues were embedded; for all of these, the “background” features were not indicative of the cue being presented.

**Figure 1:**
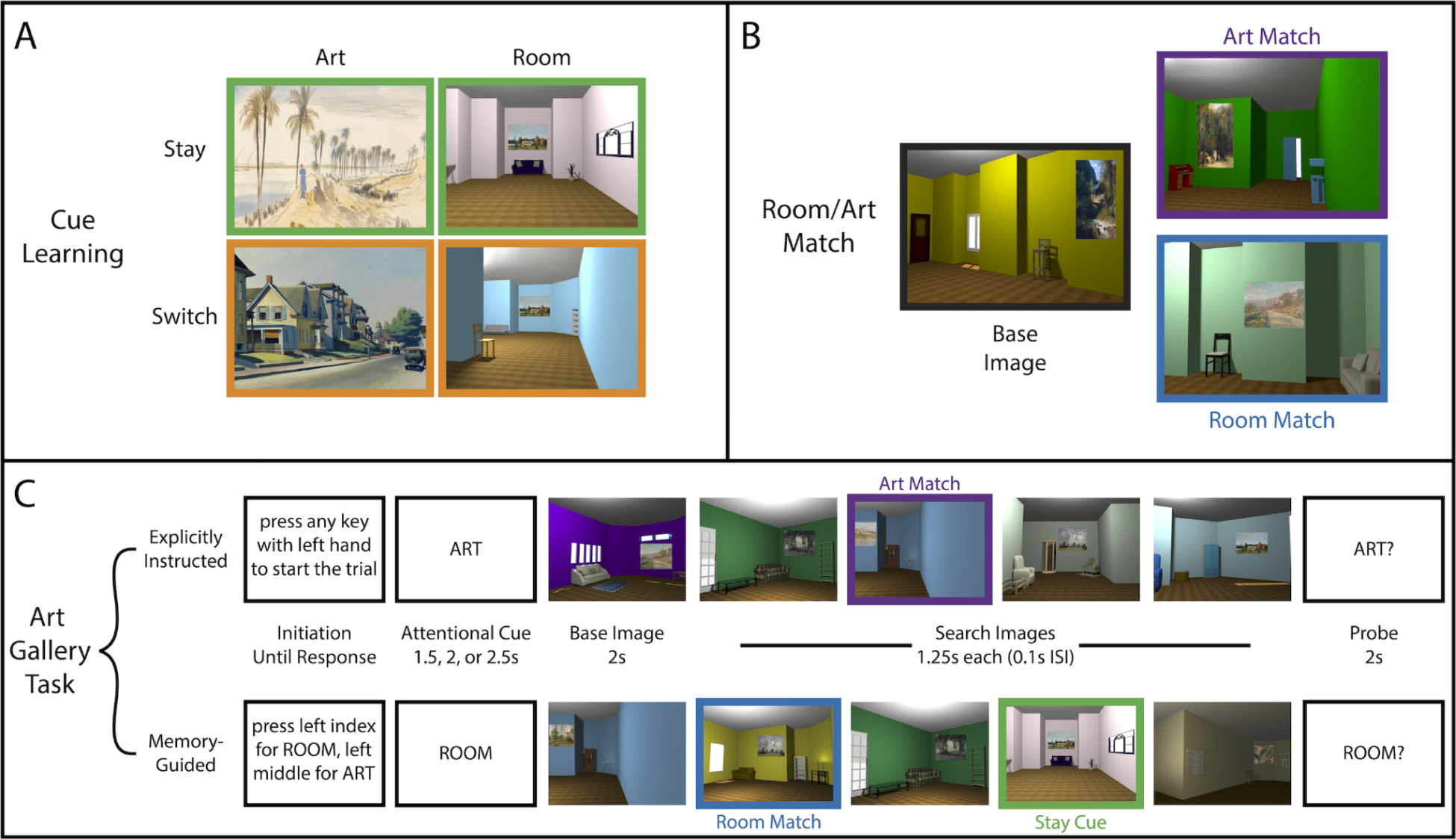
Task Overview. **(A)** Before entering the fMRI scanner, participants learned 2 “stay” cues (one painting, one room) and 2 “switch” cues (one painting, one room). These cues were embedded into the set of search images for the attention task in the fMRI scanner, and indicated how the participant should direct their attention on the following trial during the memory-guided condition. Stay cues instructed the participants to continue in the same attentional state for the following trial (i.e., art→art or room→room). Switch cues instructed the participants to change their attentional state for the following trial (i.e., art→room or room→art). Stay/switch cues were also embedded in the search images for the explicitly instructed condition but were not relevant for that task; stay/switch cues were only informative in the memory-guided condition. **(B)** For each trial of the attention task, participants saw a single “base image” of a virtual room with a painting hanging on a wall. Participants could either pay attention to the spatial layout of the room in the image *or* to the artistic style of the art on the wall. Depending on their attentional goal, participants were instructed to look for an image that either showed the same room as the base image from a different perspective (room trial) or a painting with a similar style as that in the base image (art trial). Shown are a sample base image and its room match and art match. **(C)** The attention task consisted of two conditions that occurred in different task runs: the explicitly instructed condition and the memory-guided condition. In each trial of the explicitly instructed condition, participants were randomly assigned either a room or art attentional state. Next, they were shown a base image and a series of search images. Finally, they were asked whether the search set contained an image that matched the room in the base image (same room from a different perspective, “room” probe) or the art in the base image (painting with a similar style, “art” probe). The memory-guided task had a similar structure, except that participants also had to attend to the stay or switch cue that was embedded in the search images. The stay or switch cue on a given trial was then used to select the attentional state at the start of the subsequent trial (e.g., a “stay” cue in a “room” trial, as shown above, should result in “room” being selected as the attentional state on the following trial). One-third of trials did not contain a stay or switch cue; on these “no-cue” trials, participants were free to select either art or room on the subsequent trial. Room stay/switch cues were only presented on room trials and art stay/switch cues were only presented during art trials.

The complete set of 129 main-task images (excluding the 12 stay/switch cue images) used in the fMRI task was created by permuting the 43 room images (20 rooms in 2 perspectives each and 3 “background” rooms also used with art stay/switch cues) with 43 paintings (20 artists with 2 paintings each and 3 “background” paintings also used with room stay/switch cues) such that each painting was paired with multiple rooms and each room was paired with multiple paintings.

From the set of 129 main-task images, 20 images were selected as “base images” (**Figure 1B**). Each base image was included in a set with 6 other images to form a “base set.” In addition to the base image, a base set included: a room match (another image containing a room with the same spatial layout as the room in the base image, shown from a different angle), an art match (an image containing a painting from the same artist as the painting in the base image), and 4 “distractor” images containing rooms and paintings that were neither art nor room matches to the base image, nor included any paintings or rooms used elsewhere in the base set. Base images could only be used in their own set, but art/room matches and distractors could also appear in other base sets. No single image in the base set could serve as both an art match and a room match to a single base image. For a single trial in the fMRI task, a base image was shown followed by 4 “search” images selected from the base set (art match, room match, 4 distractors) and pool of stay/switch cue images. “Stay” and “switch” cues on a given trial were always consistent with the attended dimension on that trial, i.e., art “stay” and “switch” cues only appeared on art attention trials, and room “stay” and “switch” cues only appeared on room attention trials. Each base set was used to create 5 trials for the explicitly instructed attention task and 5 trials for the memory-guided attention task; these trials differed in the images presented in the search set and the order in which they were presented. Art and room matches were independently likely to appear, such that an equal number of trials had both matches present in the search set, neither present, an art match only, or a room match only.

A second, separate set of stimuli was generated for a practice session, which occurred approximately 2 days before the scan. These practice stimuli consisted of 12 images with embedded stay/switch cues and 70 practice-task images. These 70 practice-task images were divided into 10 base sets consisting of a base image, art match, room match, and 4 distractors. Twelve novel images with embedded stay/switch cues were created using the same method described above. All practice stimuli were distinct from the fMRI task images — including the practice “stay” and “switch” cues — so that participants could learn the task while still preserving the novelty of the fMRI task stimuli.

A third, separate set of 82 stimuli was created for a practice session on the day of the scan. This practice set contained 70 novel art/room images as well as 12 images with embedded stay/switch cues. Importantly, the stay/switch cues used in this practice session were the same art and room images that would subsequently be used in the fMRI task; however, the non-attended “backgrounds” paired with each cue (3 room images and 3 paintings) were selected from the set of images used to create the 70 novel art/room stimuli for this practice session. In this way, participants could practice using the stay/switch cues that would ultimately be used in fMRI, while decoupling this practice from the unattended “backgrounds” of the stay/switch cues that would be present in the fMRI task.

### Task Overview

Participants performed an attention task on the combined art/room images described above (**Figure 1C**). On each trial, participants were first shown a base image, followed by a search set of 4 images. There were 2 potential attentional states: room attention and art attention. On a room attention trial, the participant paid attention to the spatial layout of the rooms; on an art attention trial, the participant paid attention to the artistic style of the paintings embedded in the rooms. Specifically, on a room attention trial, participants had to search the search set for a room match (an image showing a room with the same spatial layout as the base image, but from a different perspective); on an art attention trial, participants had to search the search set for an art match (an image showing a painting with a similar artistic style as the base image).

Both of these attention tasks (art and room) took place in 2 conditions: explicitly instructed attention and memory-guided attention. In the explicitly instructed condition, the participant was directed at the start of each trial to pay attention to either the room or the art with an explicit cue (“room” or “art”). Conversely, in the memory-guided condition, the participant searched for an embedded stay/switch cue — that could appear anywhere in the search set on a given trial — which indicated to them to either stay in the same attentional state in the following trial (“stay” cue), or to switch attentional states in the following trial (“switch” cue).

Both the memory-guided and explicitly instructed conditions required some amount of internal attention (e.g., to keep in mind the base image and task instructions) and external attention (e.g., to perceive the stimuli). Critically, however, one condition (memory-guided) required proportionally more internally directed attention and, therefore, proportionally less externally oriented attention, than the other (explicitly instructed). This is because only the memory-guided condition required individuals to identify and retrieve the meaning of stay/switch cues.

Critically, the stimuli were identical for the explicitly instructed and memory-guided conditions. Thus, the stay/switch cue images were also embedded in the search set for the explicitly instructed condition; they were just irrelevant to participants’ attentional state. This allowed us to compare these conditions without a confound of different stimuli.

Participants completed 4 adjacent runs (25 trials each) of the explicitly instructed condition and an additional 4 adjacent runs (25 trials each) of the memory-guided condition. The order of the conditions was counterbalanced across participants.

### Experimental Design

All stimuli and instructions were presented using Psychtoolbox for MATLAB. Each trial began with the presentation of an initiation screen. For the explicitly instructed condition, the initiation screen read “Press any key with the left hand to start the trial”. For the memory-guided condition, the initiation screen read “Press left index for Room, left middle for Art.” The structure of the rest of the trial was identical across the explicitly instructed and memory-guided conditions.

After the initiation screen, participants were shown their attentional state for the trial: “ROOM” for trials in which they were supposed to search for a room match and “ART” for trials in which they were supposed to search for an art match. This attentional state screen was presented for either 1.5s, 2s, or 2.5s (randomly assigned across trials). In the explicitly instructed condition, this attentional state was randomly assigned; however, in the memory-guided condition, the attentional state was determined by the participant’s key press during the initiation screen. On the first trial of a memory-guided run (25 trials), the participant was free to select whichever attentional state they preferred. On the following trials of the memory-guided run, participants used stay/switch cues embedded in trial N to select their attentional state in trial N+1. A “stay” cue indicated that the participant should maintain the same attentional state in the subsequent trial (e.g., if trial N were a room trial, a stay cue indicated that trial N+1 should also be a room trial). A “switch” cue indicated that the participant should choose the other attentional state on the subsequent trial (e.g., if trial N were a room trial, a “switch” cue indicated that trial N+1 should be an art trial). If a participant incorrectly selected their attentional state at the start of a memory-guided trial, the trial continued with whichever attentional state they selected. One-third of trials did not contain a stay/switch cue, and participants were told to select either attentional state on the following trial if this occurred. Participants were instructed to attempt to balance their choice of art and room attentional states following “no-cue” trials, but on average, they selected room trials more often (as reported in Günseli & Aly, 2020: average number of selected room trials = 16.828, 95% CI [16.299, 17.356]; average number of selected art trials = 14.655, 95% CI [14.144, 15.166]; t(28) = 5.90, p < 0.00001, d = 1.10, 95% CI [1.418, 2.927]).

After the initiation screen and the attentional state screen, participants were shown a base image for 2s. Depending on the attentional state (art or room), participants should either pay attention to the spatial layout of the room in the base image or to the style of the art in the base image. Next, participants saw 4 search images (selected from the trial’s base set and set of stay/switch cue images) each presented for 1.25s with a 0.1s inter-stimulus interval (ISI). Finally, 0.1s after the last search image, participants were shown a probe screen (2s) asking either “ART?” or “ROOM?” Participants responded “yes” or “no” to indicate if there was a match in the probed dimension using their right-hand index finger (match) or middle finger (no-match) on a button box. When presented with the probe “ART?” participants had to respond if any of the search images contained a painting that could have been painted by the same artist as the painting in the base image. After a “ROOM?” probe, participants responded whether or not one of the search images contained a room with the same spatial layout as the room in the base image. For 80% of trials across all runs, the probe matched the cue on the attentional state screen (e.g., if the cued attentional state was room, the probe read “ROOM?”); these were termed “valid” trials. However, 20% of trials across all runs were “invalid”, meaning the probe did not match the attentional state cue (e.g., if the cued attentional state was room, the probe read “ART?”). Comparing performance on valid vs. invalid trials allowed us to determine whether attention conferred a performance benefit (Posner, 1980; Stokes et al., 2012), and whether this performance benefit was matched across the memory-guided and explicitly instructed conditions. (Behavioral performance — including performance on valid trials only and the valid vs. invalid difference — was not different between these conditions, as noted in Günseli & Aly, 2020.)

Between trials, a blank screen was presented for a variable time (the inter-trial interval, ITI). Twenty-five ITI lengths were generated (truncated exponential, lambda = 1.5, mean = 6.66s, T = 9s) and were randomly shuffled to be used after each of the 25 trials across all 8 runs of the task.

After each run, participants were shown the percent of responses to the art/room probe that they had answered correctly. In memory-guided runs, participants were also shown the percent of stay/switch cues that they correctly identified.

The order of trials within a run was structured such that each of the 20 base images was shown once every 20 trials, no two adjacent trials could have the same base image, and no image could appear multiple times in the same trial. Matches for the probed attentional state were present in 50% of trials; independently, matches for the non-probed attentional state were present in 50% of trials. Stay/switch cues were presented in ⅔ of trials in both the explicitly instructed and memory-guided conditions (though they were only informative in the memory-guided condition). Finally, within each condition (explicitly instructed and memory-guided), 80% of trials were valid. Valid trials were equally distributed across the art/room attentional states, presence of probed match (present/absent), presence of non-probed match (present/absent), and stay/switch cue presence (stay cue/switch cue/no cue). It was not possible to perfectly equate the number of valid trials across each of these trial types for a given participant, but valid trials were balanced across these trial types across every 6 participants.

The design of the practice sessions mirrored the structure of the main experiment with the exception of the ITI lengths, which were randomly selected as either 2s or 2.5s. Additionally, for the practice sessions, participants were provided with feedback about their performance following their response to the art/room probe, and in the memory-guided condition, their selection of an attentional state.

### Procedure

The study was completed in 3 phases: an initial practice session (approximately 2 days before the fMRI scan), a second practice session (just before the scan), and the fMRI task. The initial practice session gave the participants an opportunity to learn the task and practice the explicitly instructed and memory-guided conditions with an entirely separate set of stimuli from those used in the fMRI task. Similarly, the second practice session allowed the participants to practice the task using another (mostly) separate set of stimuli from those in the subsequent fMRI task; however, we included the same stay/switch cues as those in the fMRI task to ensure that they were learned prior to the start of fMRI.

Both practice sessions began with a run of 10 trials of the explicitly instructed condition. Participants repeated this run until they answered at least 65% of the validly cued art/room probes correctly. Following practice of the explicitly instructed condition, participants entered a cue learning phase, in which they were shown each of the stay/switch cues and their meanings 4 times in a randomly shuffled order. Each image was shown for a minimum of 1 second, along with its meaning (i.e., stay or switch), and the participant had to press a button to advance to the next image. Participants were then tested on the stay/switch cues: they were shown all stay/switch cues one at a time (without their meanings) in a shuffled order. The participant had to correctly identify each of the stay/switch cues with a button press (to indicate whether the cue meant “stay” or “switch”) 5 times in a row, or else they would have to restart the cue learning sequence. Upon completion of the cue learning phase, participants performed a practice run of 10 trials of the memory-guided condition. This run was repeated until participants met two criteria: 1) they answered at least 65% of the validly cued art/room probes correctly and 2) they used stay/switch cues to correctly identify the attentional state (art or room) for the following trial on at least 80% of the trials.

After the second practice session, participants entered the scanner. The fMRI task consisted of 8 runs of 25 trials each. The explicitly instructed and memory-guided conditions each contained 100 trials (4 contiguous runs), with condition order counterbalanced across participants. Before each condition, participants completed 5 practice trials of the upcoming condition (explicitly instructed or memory-guided); practice trials were repeated until participants achieved at least 65% accuracy on the art/room probes and, in the memory-guided practice, correctly identified the attentional state (art or room) from the stay/switch cues on 80% of the trials. After each run of the memory-guided condition, participants were shown all 4 stay/switch cues (2 art images and 2 room images) and their meanings as a reminder. If on any run of the memory-guided condition, a participant was not able to correctly select the attentional state (art or room) on at least 85% of the trials, they repeated the cue learning procedure from the previous practice sessions before advancing to the next run.

### MRI Image Acquisition

All MRI scans were performed in a 3T Siemens Magnetom Prisma scanner using a 64-channel coil. Functional scans (8 task runs) used a multiband echo-planar imaging (EPI) sequence with the following settings: TR = 1.5s, echo time = 30ms, flip angle = 65°, acceleration factor = 3, voxel size = 2mm iso, phase encoding direction = posterior to anterior. 69 oblique axial slices (14° transverse to coronal) were collected in an interleaved order. Each participant also underwent a T1-weighted anatomical scan (resolution = 1mm iso) using a magnetization-prepared rapid acquisition gradient-echo sequence (MPRAGE). Field maps were collected for each participant using 69 oblique axial slices (resolution = 2mm iso).

### Regions of Interest

We focused on 4 bilateral regions of interest (ROIs): the hippocampus, the dorsal attention network (DAN), the ventral attention network (VAN), and the basal forebrain (**Figure 2**). The hippocampus was defined using the Harvard-Oxford atlas in FSL thresholded at 0.5 (Jenkinson et al. 2012). Both the DAN and VAN were defined using the procedure outlined in Jimenez et al. (2016). The DAN was defined by using FSLMATHS to create 9mm radius spheres around coordinates in the bilateral lateral frontal cortex centered at [+/- 48, 8, 35], anterior intraparietal cortex centered at [+/- 30, -58, 45], and posterior intraparietal cortex centered at [+/-46, -43, 45]. The VAN was defined by creating 9mm radius spheres around coordinates in the bilateral temporoparietal junction centered at [+/- 53, -34, 21], anterior cingulate cortex centered at [+/- 3, 20, 36], and anterior insula centered at [+/-32, 22, 4]. Finally, the basal forebrain region (specifically the medial septum and diagonal band of Broca) was defined using a probabilistic atlas from Zaborszky et al. (2021), thresholded at 0.5. All ROIs were binarized and registered to 1mm standard space.

**Figure 2:**
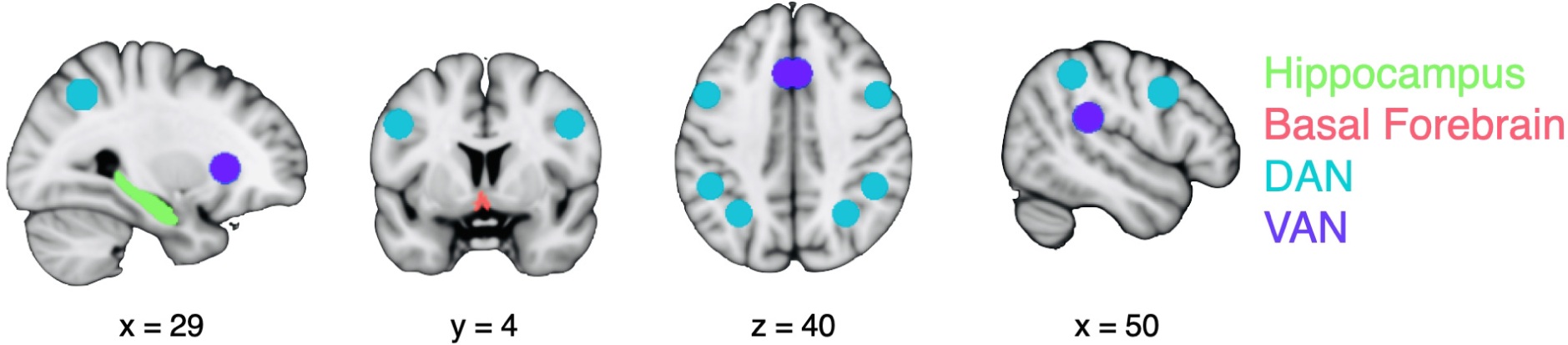
Regions of Interest. Hippocampus (green), basal forebrain (red), dorsal attention network (DAN; teal), and ventral attention network (VAN; purple) are shown on a 1mm T1 template image in MNI152 standard space. All regions of interest were bilateral.

### Preprocessing

Data were preprocessed using FEAT, FNIRT, and FSL tools (as reported in Günseli & Aly, 2020). The first 4 volumes of each functional run were removed to account for T1 equilibration; for 1 participant, only 1 volume was removed at the start of each functional run (because not enough volumes were collected at the start of each run to discard 4). Both brain extraction (BET; Smith, 2002) and motion correction (MCFLIRT; Jenkinson et al., 2002) were performed. Data were high-pass filtered (cut-off = 128s) and spatially smoothed using a 3mm full width at half maximum Gaussian kernel. Custom scripts were generated based on the FUGUE guide (https://fsl.fmrib.ox.ac.uk/fsl/fslwiki/FUGUE/Guide) to preprocess field maps. This process consisted of using the skull-stripped average of 2 magnitude images along with the phase image to generate a field map image with the fsl_prepare_fieldmap command. Field map images and magnitude images were used in FEAT preprocessing. Finally, functional images were registered to the standard MNI152 1mm anatomical image using a nonlinear warp (10mm resolution and 12 degrees of freedom). This preprocessing procedure was used in both GLMs described below.

### Statistical Analysis

#### Univariate Activity

Data were modeled using a single-trial GLM following the procedure in Günseli & Aly (2020). Each trial was made up of 3 parts (**Figure 1C**): first, an *orienting period* spanning the time from the onset of the initiation screen to the offset of the attentional state screen; second, an *image period*, spanning the 7.4s epoch from the onset of the base image until the offset of the final search image; and finally, a *probe period* spanning the 2s duration of the art/room probe screen. Each image period was modeled separately (as a 7.4s epoch); 1 additional regressor was included to model all orienting periods (modeled as epochs of varying duration depending on participant response time (RT) during the initiation screen and the length of the attentional cue); and a final regressor that modeled all probe periods (as 2s epochs). All regressors were convolved with a double-gamma hemodynamic response function as implemented in FEAT. To account for movement-related noise, 6 motion regressors were included in the model. FILM prewhitening was used to remove autocorrelation. Finally, each task run was modeled separately. All regressors of this model were registered to high-resolution 1mm MNI152 standard space.

The task included both valid trials (in which the cue and probe were the same) and invalid trials (in which participants were cued to attend to one dimension, e.g., art, but probed about the other, e.g., room). Invalid vs. valid probes lead to behavioral costs in response times and sensitivity (Aly & Turk-Browne, 2016a, b; Gunseli & Aly, 2020) and enhanced activity in the brain’s reorienting network, overlapping with both the ventral and dorsal attention networks (Aly & Turk-Browne, 2016a; Corbetta & Shulman, 2002). For these reasons, analyses of univariate activity (as well as subsequent analyses of brain-behavior correlations) focused on valid trials.

We examined univariate activity (and background connectivity, see below) for the *image period*, because this part of the task differentiates the explicitly instructed and memory-guided conditions in the proportional demand for external vs. internal attention. In the memory-guided condition, individuals must compare presented images to stay/switch cues stored in memory so that they can identify them and/or retrieve their meaning. This memory retrieval was not required in the explicitly instructed condition, for which the stay/switch cues were not relevant. We therefore expected that, during the image period, the memory-guided condition would require proportionally more internal attention and, therefore, proportionally less external attention, than the explicitly instructed condition.

For each condition and region of interest, the beta values for a given image period were first averaged across all voxels, and then separately averaged across art trials and room trials across all 4 runs of the condition. Finally, the beta values were averaged across art and room trials resulting in 1 average parameter estimate per participant per condition, equally weighted across art and room attentional states. Univariate activity was averaged across valid trials in which the participant correctly answered the subsequent art/room probe. The same pattern of results was found when using all valid trials regardless of accuracy and all trials regardless of validity or accuracy. Univariate parameter estimates were compared across the explicitly instructed and memory-guided conditions using a within-participant t-test.

#### Background Connectivity

Task-evoked fluctuations in brain activity can lead to spurious findings of functional connectivity between brain areas: if two brain regions are similarly modulated by a task, they can appear to be functionally coupled even if they are acting in isolation. For this reason, we examined functional connectivity with the “background connectivity” approach, in which correlations between brain regions are estimated after regressing out task-evoked activity (Al-Aidroos et al., 2012).

For this approach, we used a separate GLM than that used for univariate activity. Rather than modeling individual trials, the GLM was designed to capture common task-evoked activity across trials (Al-Aidroos et al., 2012). This technique of using regressors that capture the mean evoked response across trials (as opposed to single-trial regressors) is the approach taken in other studies examining background connectivity (e.g., Al-Aidroos et al., 2012; Córdova, Tompary, & Turk-Browne, 2016; Dresler et al., 2017; Li et al., 2023; Lou et al., 2015; Murty, LaBar, & Adcock, 2016; Pruitt et al., 2022; Sarpal et al., 2020; Sun et al., 2020; Tao & Rapp, 2020). Single-trial GLMs are not constrained to find common task-evoked activity across trials, and can therefore remove spontaneous activity that is not evoked by the stimuli – the critical signal for background connectivity. Thus, we followed the approach established by prior literature and used a GLM designed to capture the mean evoked response across trials.

This GLM therefore had 13 regressors that captured attentional state (art or room), trial validity (valid or invalid), participant accuracy (correct or incorrect), and trial component (orienting period, image period, probe): 1) the image periods of all valid trials with a cued (and probed) art attentional state for which participants correctly answered the probe; 2) the image periods of all valid trials with a cued (and probed) room attentional state for which participants correctly answered the probe; 3) the image periods of all invalid trials with a cued art attentional state (probed room state) for which participants correctly answered the probe; 4) the image periods of all invalid trials with a cued room attentional state (probed art state) for which participants correctly answered the probe; 5) the image periods of all valid trials with a cued (and probed) art attentional state for which participants did not correctly answer the probe; 6) the image periods of all valid trials with a cued (and probed) room attentional state for which participants did not correctly answer the probe; 7) the image periods of all invalid trials with a cued art attentional state (probed room state) for which participants did not correctly answer the probe; 8) the image periods of all invalid trials with a cued room attentional state (probed art state) for which participants did not correctly answer the probe; 9) the probe period for all valid trials for which the participant correctly answered the probe; 10) the probe periods for all invalid trials for which the participant correctly answered the probe; 11) the probe period for all valid trials for which the participant did not correctly answer the probe; 12) the probe periods for all invalid trials for which the participant did not correctly answer the probe; and 13) the orienting period of all trials. In parallel with the procedure described above for univariate activity, this GLM also included 6 motion regressors and employed FILM prewhitening. All runs were modeled separately.

The residuals of this model were registered to high-resolution 1mm MNI152 standard space. For each region of interest, the residual values were averaged across voxels and across the timepoints (5 TRs) for each image period (with onset time shifted 6s / 4 TRs from the start of the image period, to account for hemodynamic lag) resulting in a single value for each ROI for each image period. We took the approach of averaging across timepoints for each image period because this approach is functionally similar to the beta-series correlation approach (Rissman et al., 2004). This approach examines functional connectivity between brain regions based on a timecourse of beta coefficients for a trial component of interest, where the beta coefficient captures the average activity across TRs for the trial component of interest. Prior work has shown that the beta-series approach is superior to the traditional approach (in which individual timepoints are entered into the functional connectivity correlation) when there is variability in the hemodynamic response function (Cisler et al., 2014). We therefore thought this approach could help us reduce unwanted variability across brain regions that may differ in the shape of their hemodynamic response function.

For consistency with univariate analyses, the calculation of background connectivity used valid trials only, although analyses were also repeated to include invalid trials (because evoked activity was regressed out prior to conducting background connectivity analyses, trial validity should have minimal effects on background connectivity). Additionally, analyses were restricted to valid trials in which the participant correctly answered the art/room probe. This was to make sure that background connectivity was estimated across trials in which participants were in good attentional states. The same pattern of results was found when using all valid trials regardless of accuracy, all trials regardless of validity or accuracy, as well as when background connectivity was calculated without averaging across the timepoints within each trial.

For each task condition (explicitly instructed and memory-guided) and for each attentional state (room and art) the residuals for each ROI (averaged across voxels and image period timepoints) were concatenated across all aforementioned trials across all runs of the condition of interest. Background connectivity was calculated by taking the Pearson correlation of the resulting timecourses of the 2 relevant ROIs. Finally, for each condition (explicitly instructed and memory-guided) background connectivity across correct, valid art trials and across correct, valid room trials was averaged together to balance the contribution of the 2 attentional states. This procedure resulted in a single measure of background connectivity for each participant in each condition. Background connectivity was compared across the 2 conditions (explicitly instructed and memory-guided) and region pairs (basal forebrain-hippocampus and DAN-hippocampus) with a repeated-measures ANOVA on the Fisher-transformed r-values. This ANOVA also included a regressor for stay/switch cue presence (stay/switch cue present vs. absent) to allow us to test additional hypotheses about whether looking for vs. detecting stay/switch cues modulates connectivity (see Results: Background Connectivity for details). Follow-up comparisons were conducted with a within-participant t-test.

#### Temporal Multivariate-Univariate Dependence

Background connectivity provides a measure of how strongly the activity of 2 brain areas is correlated over time, but relating this measure to behavior on any given trial requires a timepoint-by-timepoint measure of functional connectivity strength. To accomplish this, Tompary et al. (2018) developed temporal multivariate-univariate dependence (TMUD) analysis to quantify the contribution of each time point to the overall correlation between 2 brain areas, giving a measure of how functional connectivity fluctuates over time. We applied the TMUD procedure to background connectivity for each task condition (explicitly instructed and memory-guided) as estimated from valid trials for both correct and incorrect judgments on the art/room probe, so that we could predict behavioral performance. First, for each region, the mean of the residual timecourse was calculated and subtracted from each individual time point in the residuals. Each mean-centered timepoint was then squared; these squared values were then summed and the square root obtained to find the “root sum-of-squares”, i.e., the square root of the sum of squared deviations from the mean. Then, each mean-centered timepoint was divided by the root sum-of-squares, resulting in a normalized timecourse of “background activity” for each region of interest. Finally, we obtained the element-wise product of 2 regions’ normalized timecourses, which provides a measure of their background connectivity strength for each trial. Indeed, the sum of these element-wise products is equal to the background connectivity between the 2 regions, offering an intuitive reason for why the element-wise product for each trial is a measure of background connectivity strength: the larger the value for a given trial, the larger the contribution of that trial to overall background connectivity.

#### Mixed Effects Logistic Regression

The relationship between brain activity and behavior was modeled using mixed-effects logistic regression. Trialwise measures of background connectivity (TMUD values calculated according to the procedure above) between the DAN and the hippocampus, and between the basal forebrain and the hippocampus, were used to predict participants’ accuracy (0 or 1) on the art/room probe for each valid trial. First, the TMUD values were z-scored; outliers were then removed on an individual participant basis using Rosner’s test for outliers as implemented in R. The model also included the attentional state of the given trial (art or room, effect coded) as well as a random intercept for each participant.

We first modeled the explicitly instructed and memory-guided conditions in the same model (effect coding the conditions), and incorporated an interaction between condition and TMUD, as follows: Accuracy ∼ TMUD + Condition + TMUD*Condition + Attentional State + (1|Participant). Separate models were run for DAN-hippocampus and basal forebrain-hippocampus.

We also ran follow-up models separately for the two conditions, as follows: Accuracy ∼ TMUD + Attentional State + (1|Participant).

Finally, an additional model for each condition was also run using univariate activity in the DAN in place of TMUD values.

#### Whole-Brain Analysis

Finally, we conducted a whole-brain background connectivity analysis analogous to our ROI approach. We looked for voxels whose residual activity timecourse was correlated with the residual activity timecourse in the hippocampus during the image period, using the same trials as those used for the ROI analysis (valid trials in which the participant responded correctly on the art/room probe). We did this for both the explicitly instructed and memory-guided conditions, and then performed group-level analyses on the condition differences using FSL’s “randomise” function.

## Results

### Validation of Background Connectivity Approach

Prior to examining our hypotheses regarding background connectivity, we first sought to ensure that we effectively removed task-evoked activity with our GLM. To that end, we examined the residuals from the background connectivity GLM (described in **Methods**), averaging across timepoints separately for art and room trials. Our prior studies have found robust differences in univariate activity across art and room trials in the hippocampus and across the brain more generally (Aly & Turk-Browne, 2016a, 2016b). We verified that we effectively removed this task-evoked activity in our background connectivity GLMs, for the trials of interest used in subsequent analyses (i.e., valid trials with correct responses for the art/room probe). Averaging the residuals for these trials confirmed no difference between art and room attentional states in the hippocampus (memory-guided: t(28) = -0.50, p = 0.62; explicitly instructed: t(28) = 0.41, p = 0.68), basal forebrain (memory-guided: t(28) = 0.53, p = 0.60; explicitly instructed: t(28) = -1.63, p = 0.11), DAN (memory-guided: t(28) = -0.29, p = 0.77; explicitly instructed: t(28) = -1.28, p = 0.21), or a control region of interest, the ventral attention network (VAN; memory-guided: t(28) = -0.11, p = 0.92; explicitly instructed: t(28) = -0.57, p = 0.57). There were also no differences between the averaged residuals for the memory-guided and explicitly instructed conditions in any region of interest (hippocampus: t(28) = 1.52, p = 0.14; basal forebrain: t(28) = 0.24, p = 0.81; DAN: t(28) = -0.23, p = 0.82; VAN: t(28) = 1.46, p = 0.15). Thus, these residuals — to be examined in our background connectivity analyses — did not contain reliable information about attentional states or task conditions.

### Background Connectivity

Our main analyses centered on changes in hippocampal network interactions across the explicitly instructed and memory-guided attention conditions. We hypothesized that we would find stronger interactions between the basal forebrain and the hippocampus during explicitly instructed vs. memory-guided attention. We expected the opposite effect for DAN and hippocampus, with higher background connectivity during memory-guided vs. explicitly instructed attention.

To test these hypotheses, we measured “background connectivity” by calculating functional connectivity between the regions of interest after removing task-evoked activity (Al-Aidroos et al., 2012). We then conducted an analysis of variance to determine if background connectivity between our regions of interest varied by condition (memory-guided vs. explicitly instructed).

We also included a regressor for whether the trial included a stay or switch cue (cue present) vs. not (cue absent). If hippocampal connectivity flips between the basal forebrain and DAN specifically when a stay/switch cue is encountered in the memory-guided condition (but not in the explicitly instructed condition, for which these cues are not relevant for behavior) then we should observe a region pair (hippocampus-basal forebrain vs. hippocampus-DAN) by condition (explicitly instructed vs. memory-guided) by cue presence (present vs. absent) interaction. Alternatively, if the hippocampal connectivity flip is triggered by the need to compare presented images to the stay/switch cues in memory, whether the stay/switch cues ultimately appear or not, then no three-way interaction would occur.

We observed the hypothesized interaction between condition (memory-guided vs. explicitly instructed) and region pair (basal forebrain-hippocampus vs. DAN-hippocampus), F(1, 28) = 10.19, p = 0.003, generalized η^2^ = 0.03 (**Figure 3A**). Given this interaction, we conducted follow-up t-tests to compare background connectivity patterns across conditions. Consistent with our hypothesis, background connectivity between the basal forebrain and the hippocampus was significantly stronger during explicitly instructed vs. memory-guided attention, t(28) = 3.03, p = 0.005, 95% CI [0.048, 0.250], Cohen’s dz = 0.56 (**Figure 3A**, red). Conversely, background connectivity between the DAN and hippocampus was significantly higher during memory-guided vs. explicitly instructed attention, t(28) = -2.24, p = 0.03, 95% CI [-0.215, -0.01], Cohen’s dz = 0.42 (**Figure 3A**, teal).

**Figure 3:**
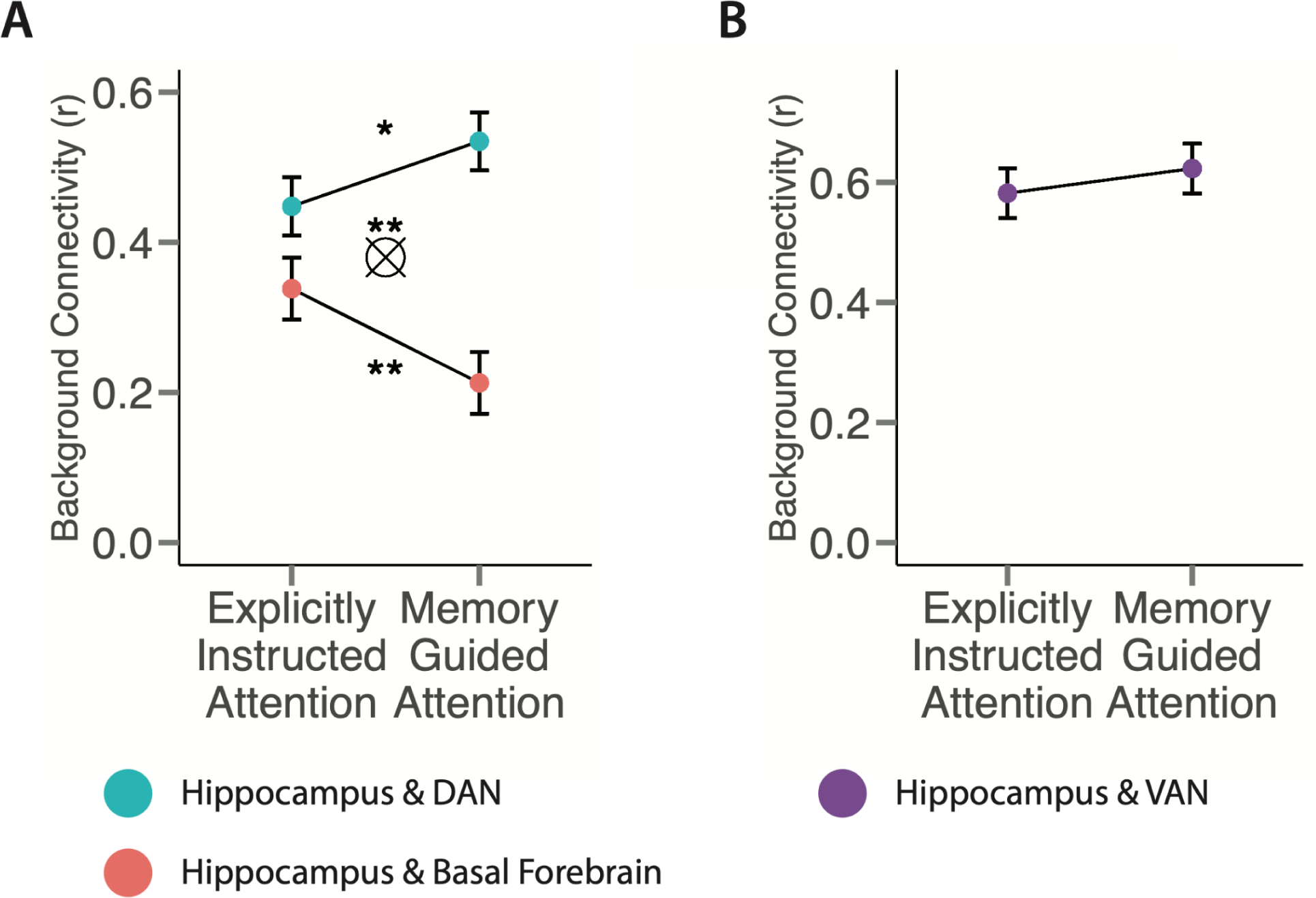
Background connectivity in regions of interest. **A.** Background connectivity was calculated between the hippocampus and the dorsal attention network (DAN) and between the hippocampus and basal forebrain for each attention condition (explicitly instructed and memory-guided). Solid circles represent the mean background connectivity across participants in each condition, and error bars represent the standard error of the within-participant difference between conditions (separately for hippocampus-DAN and hippocampus-basal forebrain connectivity). Background connectivity between the DAN and the hippocampus was significantly higher during the memory-guided vs. explicitly instructed condition. Conversely, background connectivity between the basal forebrain and hippocampus was significantly higher in the explicitly instructed vs. memory-guided condition. The X symbol represents a significant interaction between attention condition and inter-regional interactions. **B.** Background connectivity between the VAN and the hippocampus was not significantly different across conditions. (* p < .05, ** p < .01, *** p < .001).

Turning to stay/switch cue presence, we found no main effect of, nor interactions involving, cue-presence (all *p*s > 0.1). The lack of a three-way interaction between condition, region pair, and stay/switch cue presence suggests that the requirement to compare presented images to stay/switch cues stored in memory may modulate hippocampal connectivity between the DAN and basal forebrain, whether or not a stay/switch cue is ultimately presented. An alternative, however, is that we were not well-powered to detect a three-way interaction — but a follow-up exploratory two-way ANOVA for the memory-guided condition only also failed to yield an interaction between cue presence and region pair, F(1, 28) = 0.02, p = 0.90, generalized η^2^ = 0.00007. This null result is consistent with the interpretation that DAN-hippocampal connectivity may index the allocation of internal attention toward stay/switch cues stored in memory, so that these internal representations can be compared to externally presented stimuli. Whether a match is ultimately presented or not may matter less than the need to allocate attention to stored representations. We return to this issue further in the Discussion. Because there was no main effect of, nor interactions involving, cue presence, subsequent analyses included both types of trials; however, when significant effects were observed, follow-up analyses were conducted to ensure that the effects hold when analyzing cue-present trials only.

The interaction between region pair (hippocampus-DAN, hippocampus-basal forebrain) and attention condition (memory-guided, explicitly instructed) remained significant when using all valid trials regardless of accuracy (F(1, 28) = 13.34, p = 0.001, generalized η^2^ = 0.02), all trials regardless of validity or accuracy (F(1, 28) = 17.77, p = 0.0002, generalized η^2^ = 0.03), as well as when background connectivity was calculated without averaging across the timepoints within each trial (F(1, 28) = 7.57, p = 0.01, generalized η^2^ = 0.02).

We next tested whether the switches in hippocampal background connectivity, reported above, were relatively selective to our networks of interest. To that end, we examined hippocampal background connectivity with the ventral attention network (VAN). The VAN is associated with bottom-up attention and shows increased activity for salient stimuli (Corbetta and Shulman, 2002; Vossel et al., 2014). Although the VAN is centrally involved in attentional orienting, we did not expect this network to differentially interact with the hippocampus across the explicitly instructed and memory-guided attention conditions.

The logic for this hypothesis was as follows. To keep stimuli identical across tasks, the stay/switch cues were embedded in both memory-guided and explicitly instructed trials; however, in the latter, they were irrelevant to the attentional state on the following trial. Because participants were extensively trained, prior to the fMRI task, to identify stay/switch cues, the stay/switch cues may have captured bottom-up attention to some extent in both the memory-guided and explicitly instructed conditions. If so, then the VAN — involved in bottom-up attention capture, as noted above — may not show differential background connectivity with the hippocampus across tasks.

We first tested whether the VAN shows differential univariate activity across conditions. As we expected, we observed no difference, t(28) = 0.15, p = 0.88, 95% CI [-3.84 4.44], Cohen’s dz = 0.03. Furthermore, as we hypothesized, background connectivity between the VAN and the hippocampus was not significantly different across the explicitly instructed and memory-guided conditions, t(28) = -0.96, p = 0.35, 95% CI [-.19, 0.07], Cohen’s dz = -0.18 (**Figure 3B**). Thus, switches in hippocampal background connectivity across conditions did not extend more broadly to other attention networks in the brain.

### Relationship between Background Connectivity and Behavior

We next sought to test whether the strength of background connectivity between the hippocampus and our regions of interest (DAN, basal forebrain) was related to performance on the attention task. To test this, we first used a TMUD analysis to generate trial-by-trial measures of background connectivity. We then used logistic regression to model the relationship between these trial-wise background connectivity measures and participants’ accuracy on the art/room probe. We also included attentional state (art or room) as a control regressor in the model, to account for differences in accuracy across these attention tasks. Thus, we predicted trial-wise accuracy based on background connectivity (with separate models for DAN-hippocampus and basal forebrain-hippocampus), condition (memory-guided, explicitly instructed), attentional state (art, room), and the interaction between background connectivity and condition.

We first examined the relationship between DAN-hippocampus background connectivity and behavior. We found a significant interaction between trial-wise background connectivity strength and attention condition (memory-guided or explicitly instructed), β = 0.32, SE = 0.13, p = 0.02 (**Figure 4A-C**). To probe this interaction, we conducted separate logistic regression models for the two conditions. There was a significant positive relationship between DAN-hippocampus background connectivity and art/room probe accuracy in the memory-guided condition, β = 0.26, SE = 0.09, p = 0.004. There was no relationship between DAN-hippocampus background connectivity and probe accuracy in the explicitly instructed condition, β = -0.06, SE = 0.10, p = 0.56. We observed the same pattern of results when restricting the analysis to only those trials that contained a stay/switch cue, i.e., excluding no-cue trials (memory-guided: β = 0.35, SE = 0.11, p = 0.002; explicitly instructed: β = 0.04, SE = 0.11, p = 0.72).

**Figure 4:**
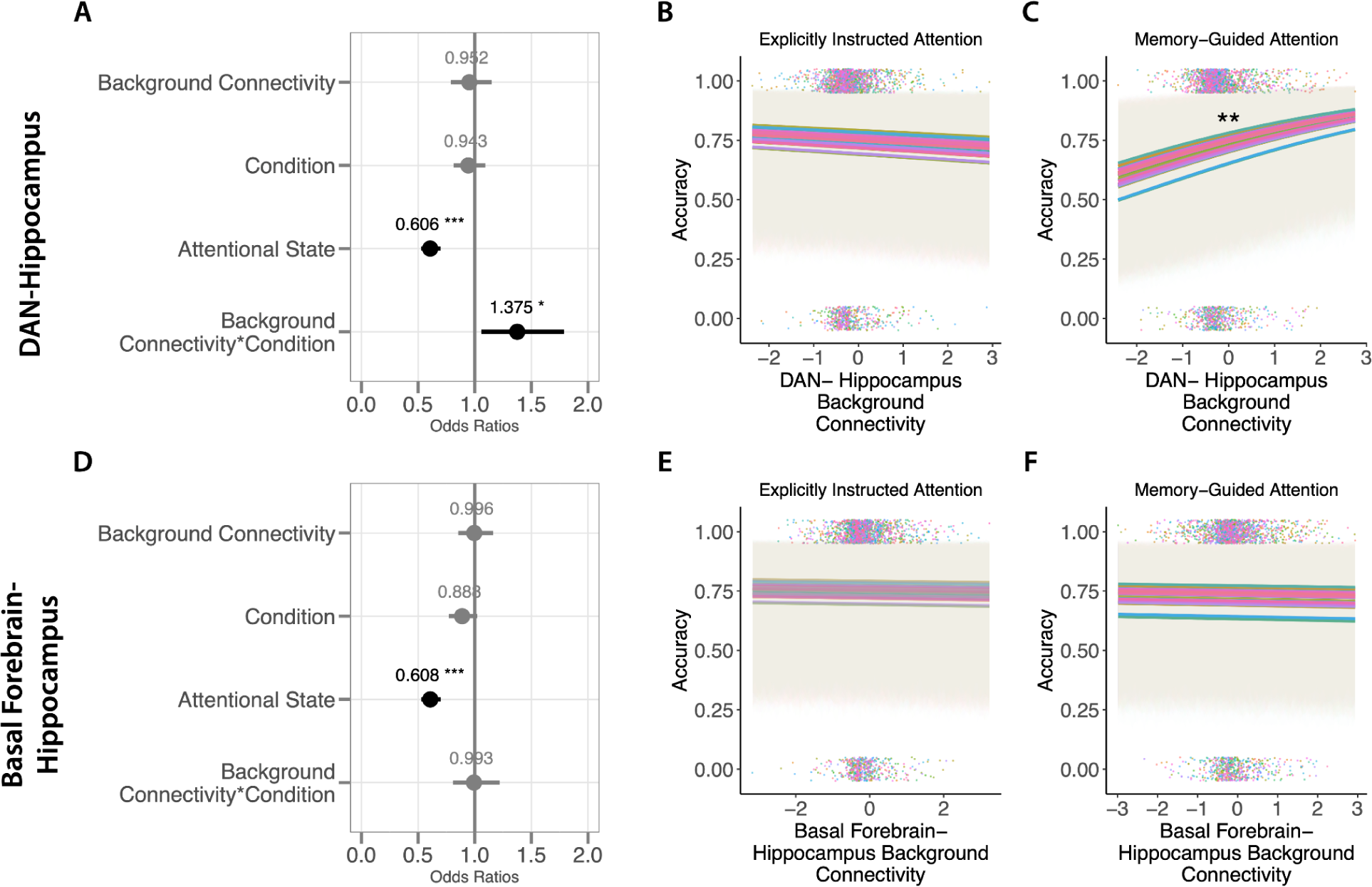
Relationship between DAN-hippocampus (top) and basal forebrain-hippocampus (bottom) background connectivity and accuracy on the attention task. **Top: (A)** The parameter estimates of the full model are displayed as odds ratios; background connectivity = trial-wise z-scored DAN-hippocampus background connectivity; condition = memory-guided or explicitly instructed (effect coded); attentional state = art or room (effect coded). DAN-hippocampus background connectivity did not predict accuracy on the attention task in the explicitly instructed condition **(B)** but did predict accuracy in the memory-guided condition **(C)**. **Bottom:** The parameter estimates of the full model are depicted as odds ratios **(D)** with the same regressors as in **(A)** except with basal-forebrain hippocampus background connectivity. Basal forebrain-hippocampus background connectivity did not predict attention task performance in either the explicitly instructed **(E)** or memory-guided **(F)** conditions. Colored lines represent the fitted model for each participant and colored dots represent trial-wise connectivity for each participant separately for incorrect (0) and correct (1) trials. (* p < .05, ** p < .01, *** p < .001).

Basal-forebrain hippocampal background connectivity strength did not predict accuracy on the art/room probe (β = -0.004, SE = 0.08, p = 0.96) nor was there an interaction between background connectivity strength and condition (β = -0.01, SE = 0.11, p = 0.95). Thus, basal-forebrain hippocampus background connectivity generally did not predict behavior, nor was there evidence for differential effects across the memory-guided (β = -0.01, SE = 0.07, p = 0.85) and explicitly instructed (β = -0.01, SE = 0.08, p = 0.89) conditions (**Figure 4D-F**).

### Analysis of Univariate Activity

We found that DAN-hippocampal background connectivity predicted behavioral performance on the art/room probe in the memory-guided condition. Are these results selective to DAN-hippocampal background connectivity or does DAN activity on its own predict performance? It is possible, for example, that the DAN is more engaged in the memory-guided vs. explicitly instructed condition because the former requires more top-down control and multitasking (Majerus et al., 2012; Majerus et al., 2018). To test this, we first probed whether task-evoked univariate DAN activity was higher during memory-guided attention than explicitly instructed attention; however, there was no difference between conditions, t(28) = 1.47, p = 0.15, 95% CI [-1.66 10.10], Cohen’s dz = 0.27 **(Figure 5A**). Thus, the memory-guided condition may not have required more top-down attentional effort than the explicitly instructed condition, perhaps because participants were extensively trained on the task; indeed, this is consistent with the lack of a behavioral difference between conditions (as reported in Günseli & Aly, 2020).

**Figure 5:**
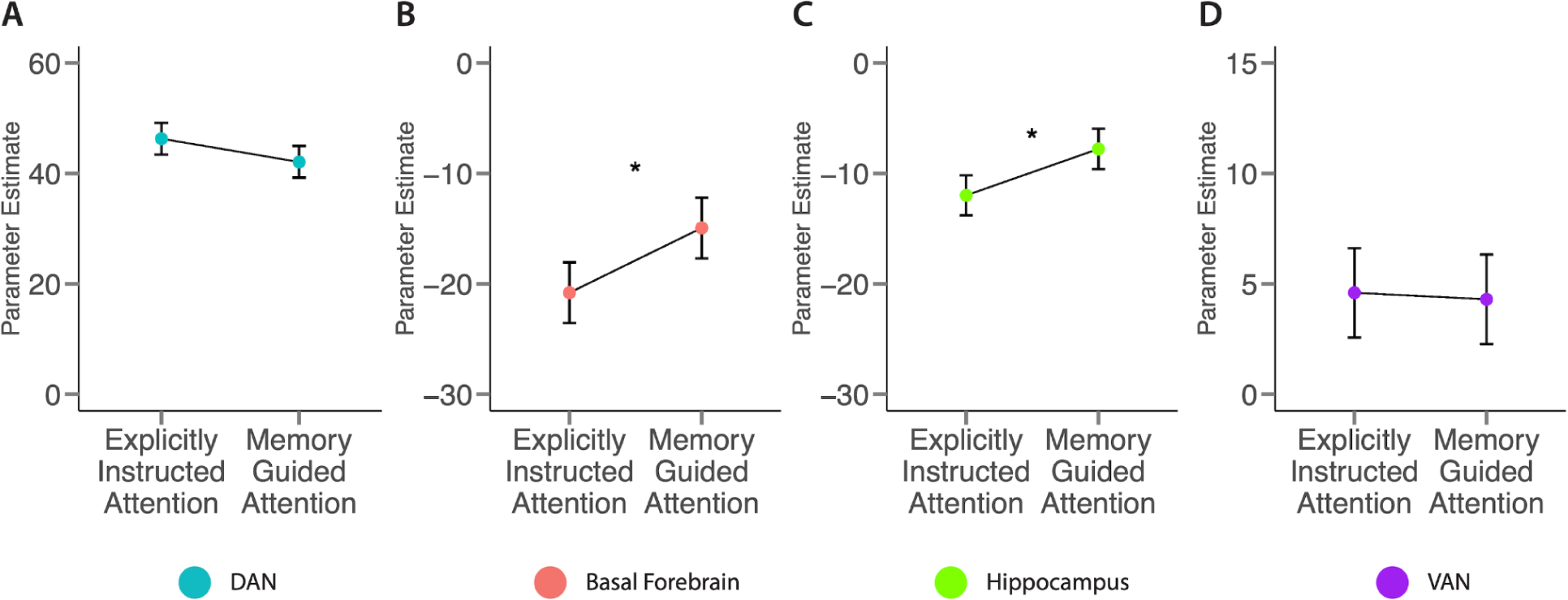
Univariate activity. Univariate activity in our regions of interest. **(A)** Dorsal attention network. **(B)** Basal forebrain. **(C)** Hippocampus. **(D)** Ventral attention network. Circles represent the average parameter estimates across participants and error bars represent the standard error of the within-participant difference between conditions. (* p < 0.05)

Although there was no difference in DAN activity across conditions, it is possible that DAN activity may predict behavior in the memory-guided but not explicitly instructed condition. DAN activity did not, however, predict art/room probe accuracy in either condition (memory-guided: β = 0.03, SE = 0.05, p = 0.50, explicitly instructed: β = -0.03, SE = 0.06, p = 0.62). Thus, DAN-hippocampus background connectivity, but not DAN univariate activity, predicts behavioral performance in the memory-guided condition.

For completeness, we also examined univariate activity in our other regions of interest (**Figure 5B-D**; note that the VAN result was reported earlier in Results: Background Connectivity). Basal forebrain univariate activity was significantly higher in the memory-guided vs. explicitly instructed condition, (t(28) = -2.14, p = 0.04, 95% CI [-11.49 -0.24], Cohen’s dz = 0.40; **Figure 5B**). Similarly, hippocampal univariate activity was significantly higher in the memory-guided vs. explicitly instructed condition, (t(28) = -2.32, p = 0.03, 95% CI [-7.92 -0.49], Cohen’s dz = 0.43; **Figure 5C**) (note that this analysis replicates the finding originally reported in Günseli & Aly, 2020, with different trials). Thus, despite individually showing higher task-evoked activity during memory-guided vs. explicitly instructed attention, hippocampus and basal forebrain show greater background connectivity with one another during explicitly instructed vs. memory-guided attention. This suggests that task-evoked univariate activity and background connectivity yield complementary insights into brain regions’ roles in memory-guided attention.

Together, our results show that the hippocampus exhibits a flip in background connectivity between the basal forebrain and DAN as proportional demands on external vs. internal attentional goals vary. Furthermore, hippocampal coupling with the DAN is associated with better attentional performance in the memory-guided vs. explicitly instructed condition. These results were specific to our networks and measures of interest, and did not extend to hippocampal coupling with the VAN or to univariate DAN activity.

### Exploratory Analysis of the mPFC

The cholinergic basal forebrain is a small neuromodulatory structure that is not typically considered a source of top-down attentional control. This leads to the question of which brain areas may be driving the basal forebrain to in turn modulate the hippocampus. One possibility is the medial prefrontal cortex (mPFC), which is connected to both the cholinergic basal forebrain (Gaykema et al., 1991; Markello et al., 2018) and the hippocampus (Dolleman-van der Weel et al., 2019; Hyman et al., 2005). To examine mPFC involvement in our task, we conducted an exploratory analysis by defining 9 mm sphere ROIs around peak voxels in medial prefrontal areas that show connectivity with the medial septum and diagonal band of Broca (Markello et al., 2018), the areas of the cholinergic basal forebrain that project to hippocampus, and which we examined in the current study. We found that this mPFC ROI was not differentially coupled with the hippocampus across conditions, t(28) = -0.56, p = 0.58, 95% CI [-0.09 0.16], Cohen’s dz = -0.10, but it was marginally more coupled with the basal forebrain during the explicitly instructed vs. memory-guided condition, t(28) = 1.77, p = 0.09, 95% CI [-0.02 0.23], Cohen’s dz = 0.33. This exploratory analysis therefore offers weak evidence consistent with a proposed role for the mPFC in driving the cholinergic basal forebrain during externally oriented attention. It also raises the possibility that some mPFC connectivity with the hippocampus may be independent of external vs. internal attentional states, with mPFC exerting additional effects on the hippocampus via the basal forebrain to drive externally oriented attention.

### Whole-Brain Analysis

Finally, we conducted a whole-brain background connectivity analysis to look for voxels whose residual activity timecourse was differentially correlated with that of the hippocampus across the memory-guided and explicitly instructed conditions. This analysis served two purposes. First, the basal forebrain is a small region that can be difficult to image (Zaborszky et al. 2008); thus, a whole-brain analysis allowed us to assess whether our basal forebrain-hippocampal coupling effect was centered on the basal forebrain ROI or was more sparse and noisy over a larger area. Second, this analysis allowed us to examine whether there were robust effects outside of our networks of interest.

We first examined the explicitly instructed > memory-guided contrast, with a focus on the basal forebrain. We applied a liberal threshold of uncorrected p < .01 to visualize the pattern of effects around the basal forebrain without statistical corrections. This allowed us to determine if our effect was relatively localized or if there was a more distributed and noisy signal that happened to overlap with the basal forebrain. We observed a cluster that largely overlapped with our basal forebrain ROI, suggesting that our targeted region-of-interest approach was effective at capturing a robust basal forebrain signal – a signal that was not sparse and expansive over neighboring regions (**Figure 6**). Note that the mPFC clusters in this liberally thresholded analysis are broadly consistent with our proposal (mentioned above) that a circuit linking mPFC, the basal forebrain, and hippocampus may play a role in prioritizing externally vs internally oriented attention.

**Figure 6:**
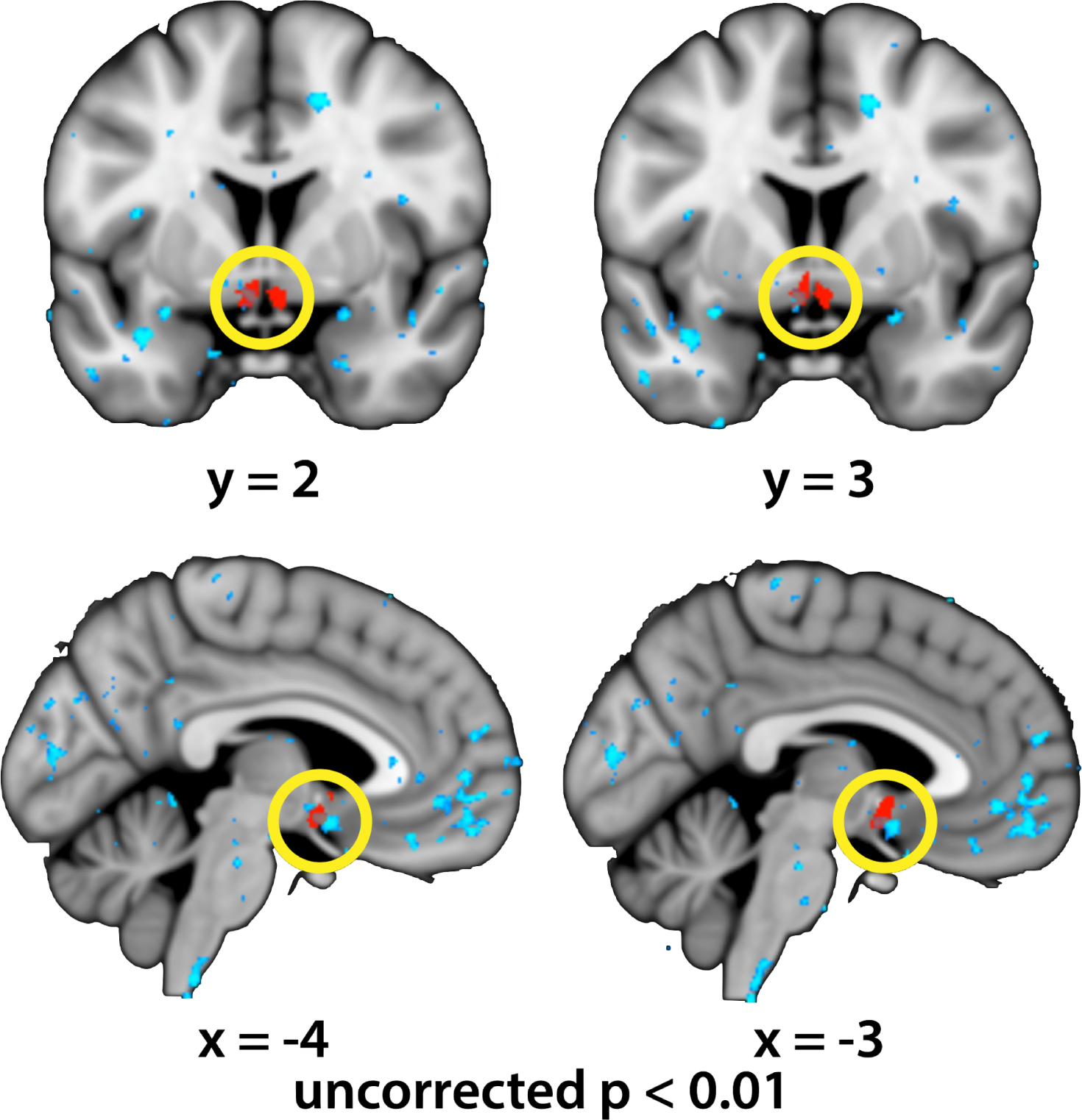
Basal forebrain-hippocampal connectivity from a whole-brain analysis. Explicitly instructed > memory-guided contrast, with a liberal threshold of uncorrected p < .01. The basal forebrain ROI is shown in red and the statistical map is shown in blue; both are slightly translucent to enable overlap to be detected. The whole-brain analysis revealed a cluster that largely overlapped with the basal forebrain ROI, as opposed to a noisier or sparser signal across a broad region.

We then applied statistical corrections to determine whether there were reliable condition differences in hippocampal background connectivity across the brain (family-wise error corrected p < .05). No clusters emerged in this whole-brain analysis for either contrast (memory-guided > explicitly instructed or vice versa). Thus, our targeted region-of-interest approach was effective at identifying hippocampal network switches and we do not seem to have missed robust effects elsewhere — although additional analyses with more ROIs beyond the ones used here may also yield effects.

## Discussion

Everyday experience requires individuals to constantly switch between attending to the external environment and their internal memories and thoughts. How is this attentional switching coordinated? We focused on hippocampal functional connectivity profiles with attention networks in the brain, given evidence that the hippocampus is involved in both internal and external attention (Eichenbaum, 2004; Aly & Turk-Browne, 2016a, 2016b; Honey et al., 2017; Córdova et al., 2019; Ruiz et al., 2020). We compared background connectivity patterns across two tasks that varied in the proportional amounts of internal vs. external attention that they demanded: the memory-guided task required proportionally more internal attention, and proportionally less external attention, than the explicitly instructed task. Consistent with our hypothesis, we found that the memory-guided task was associated with both significantly higher DAN-hippocampus connectivity and significantly lower basal forebrain-hippocampus connectivity than the explicitly instructed task. Importantly, these changes in hippocampal background connectivity did not extend to the ventral attention network (VAN), a region we hypothesized would be involved in both tasks due to its role in bottom-up attention (Corbetta and Shulman, 2002; Vossel et al., 2014). Finally, hippocampal coupling with the DAN predicted attentional behavior on the memory-guided but not explicitly instructed task.

We propose that DAN-hippocampus background connectivity may allow for the efficient retrieval of stay/switch cue memories in the memory-guided attention task, thus freeing up attentional resources to process external visual signals and perform accurately on art/room match detection. DAN-hippocampus background connectivity may not have been as important in predicting task performance in the explicitly instructed condition because that task placed fewer demands on long-term memory retrieval. Unexpectedly, however, there was no relationship between hippocampal-basal forebrain coupling and attentional behavior. This was surprising because the basal forebrain should bias the hippocampus toward an externally oriented state (Newman et al., 2012; Tarder-Stoll, Jayakumar, et al., 2020; Decker & Duncan, 2020), which should in turn improve externally oriented attentional behavior. We return to this null effect below when we discuss limitations.

Together, our results suggest that switches between internal and external attention may be coordinated at least in part by changes in how different attention networks communicate with the hippocampus. This pattern of results raises an important question about the causal relationship between our regions of interest. fMRI does not allow us to determine which regions are the operators and which are the targets; however, based on past work, we would expect that that the basal forebrain operates on the hippocampus via cholinergic modulation of external vs. internal input strengths (Hasselmo et al., 2006; Newman, et al., 2012; Tarder-Stoll, Jayakumar, et al., 2020; Decker & Duncan, 2020). The basal forebrain, in turn, may be driven by the medial prefrontal cortex (Gaykema et al., 1991). Separately, we hypothesize that the DAN may drive the hippocampal switch to internal attention by allocating attentional resources that facilitate the ability of the hippocampus to access internal representations (Cabeza, 2008; Cabeza et al., 2008; Wagner et al., 2005).

### Relation to existing work

The relationship between the basal forebrain and the hippocampus has been a rich area of research within the computational modeling and rodent literatures (Hasselmo, 1995; Parent & Baxter, 2004; Newman et al., 2012; Honey et al., 2017). Within the hippocampus, acetylcholine from the basal forebrain strengthens afferent (external) synapses and suppresses reciprocal (internal) synapses (Hasselmo 2006; Tarder-Stoll, Jayakumar, et al., 2020; Decker & Duncan, 2020). Together, this modulation of external and internal connections can drive an externally oriented state in the hippocampus. Our findings critically expand on this work by providing empirical evidence in humans that functional connectivity between the basal forebrain and the hippocampus is strengthened when there are greater demands for external attention. Our results therefore suggest that top-down goals to attend to the external world may modulate coupling between the cholinergic basal forebrain and hippocampus, driving the hippocampus toward an externally oriented state.

Recent work by Li et al. (2022) explored external and internal states across the brain using background connectivity, the same approach used in the current study. Li et al. found that sustained background connectivity within the default mode network characterized an internally oriented state, whereas stable patterns of background connectivity within the DAN/cognitive control network defined an externally oriented state. Moreover, the retrosplenial cortex flexibly switched between these two networks during internal vs. external attention. This work, along with other research (Spreng et al., 2010; Kim, 2015), emphasizes the role of the DAN as an externally oriented, task-positive network; however, Li et al. argue that this externally oriented state does not preclude situations in which the DAN is recruited for internal attention as well. Furthermore, they propose that the hippocampus (which did not emerge in their whole-brain analysis) may shift between externally and internally oriented modes (Li et al., 2022).

Although Li et al. did not find significant shifts in network connectivity with the hippocampus, we would still expect our results to replicate in future experiments — with the key caveat that the attentional task must be shown to modulate the hippocampus and tax relational and/or spatial attention. Indeed, we selected this task because it robustly recruits the hippocampus (Aly & Turk-Browne, 2016a; Aly & Turk-Browne, 2016b; Günseli & Aly, 2020; Ruiz et al., 2020). A task design that does not leverage relational or spatial attention may not modulate hippocampal activity, as we have argued in prior work (Aly & Turk-Browne, 2016a; Aly & Turk-Browne, 2017; Aly & Turk-Browne, 2018; Córdova et al., 2019). Indeed, based on our past work (Aly & Turk-Browne, 2016a; Aly & Turk-Browne, 2016b; Aly & Turk-Browne, 2018; Günseli & Aly, 2020; Ruiz et al., 2020), we expect the hippocampus to be most involved in attention tasks that require processing of higher-order relationships between different items that are distinct in their low-level features, e.g., rooms with the same spatial layout from different perspectives, with different wall colors and furniture exemplars. Thus, future studies looking to examine hippocampal network switches as a function of internal vs. external attention demands should ensure that the tasks used reliably engage the hippocampus.

As hypothesized, we found that the hippocampus exhibits switches in background connectivity patterns in tasks that are known to reliably engage the hippocampus and vary in external vs. internal attention requirements; however, contrary to what may be expected based on Li et al., we found that the DAN is more strongly coupled with the hippocampus during attention tasks with proportionally greater demands on *internal* attention. Because the DAN is involved in both external attention (e.g., Li et al.) and internal attention (e.g., Stokes et al., 2012), it is possible that the recruitment of the DAN to help shift the hippocampus towards internal processing is dependent on the particular cognitive load and the demands of the given task. Future research can systematically compare DAN-hippocampus functional connectivity in various attention tasks to determine when and how the DAN may prioritize internally vs. externally oriented attention.

Based on our results, we would expect that DAN-hippocampal functional connectivity is particularly important when there is a demand to allocate attentional resources toward memory retrieval in the midst of another ongoing task (Cabeza et al., 2008). We would also expect that the demand to allocate internal attention to memory retrieval is more important than the demand to successfully identify a previously encountered item in the environment. In particular, we found that hippocampal-DAN connectivity in the memory-guided condition did not vary based on whether a stay/switch cue was encountered or not. This indicates that DAN-hippocampal connectivity may index the allocation of internal attention toward stay/switch cues stored in memory, so that these internal representations can be compared to externally presented stimuli. Whether there is ultimately a match in the environment to these internal representations or not may matter less than the need to maintain these internal representations in the first place. This proposal, however, should be tested in future work because our study was not designed to test this specific hypothesis.

Finally, our choice of regions of interest for the hippocampal connectivity analysis was motivated by prior work linking the cholinergic basal forebrain and dorsal attention network to external and internal states, respectively (Hasselmo, 2006; Cabeza, 2008; Cabeza et al., 2008). Although theoretically motivated, one limitation of our approach is that the basal forebrain and DAN are very different structures with different modes of operation. The basal forebrain is a small neuromodulatory structure whereas the dorsal attention network consists of several frontoparietal areas involved in top-down attentional control. In this way, these regions may instantiate different types of information processing. Although those differences did not prevent us from detecting the hypothesized interaction in hippocampal connectivity, future work can extend our findings by establishing which prefrontal cortical areas modulate the basal forebrain to in turn drive the hippocampus. Based on past work examining structural and functional connectivity of the basal forebrain (Gaykema et al., 1991; Markello et al., 2018), we think a likely candidate is the medial prefrontal cortex (mPFC) – although an exploratory analysis with our data only provided weak evidence for this hypothesis (see Results: Exploratory Analysis of the mPFC). Future work can test when and how mPFC drives the basal forebrain to modulate the hippocampus, and can likewise determine which neuromodulatory systems may enable the DAN to push the hippocampus into an internally oriented mode. Such studies would be able to establish the circuit mechanisms by which specific prefrontal regions may modulate the hippocampus to switch between internal and external attention, via action of potentially distinct neurotransmitter systems.

### Limitations and Future Directions

We found that DAN-hippocampus background connectivity during the memory-guided condition (but not the explicitly instructed condition) predicted performance on art/room match detection. This null result for the explicitly instructed condition is intriguing because one might have instead predicted a *negative* relationship. Because long-term memory retrieval demands were minimal for the explicitly instructed condition, moments of increased DAN-hippocampus background connectivity in this condition might instead reflect internally oriented distraction (e.g., mind wandering) which should then hamper performance. Future research may uncover such a negative relationship between DAN-hippocampus functional connectivity and performance on external attention tasks, perhaps in settings in which mind wandering might be particularly common.

In contrast, because internal attention demands were higher for the memory-guided condition, we expected DAN-hippocampal background connectivity to benefit attentional performance — and indeed observed such an effect. This relationship may reflect the importance of DAN-hippocampus background connectivity in retrieving memories of the stay/switch cues in the memory-guided condition, which in turn frees up cognitive resources for performing the primary art/room match detection task. If this is true, one might also predict that stronger background connectivity between the DAN and hippocampus might be associated with better performance in selecting the correct attentional state in the memory-guided condition (based on the stay/switch cues). Our study was not ideally set up to test this particular hypothesis, however, for two reasons. First, accuracy in selecting attentional states in the memory-guided condition approached ceiling performance (94.9% for stay cues and 96.7% for switch cues, as reported in Günseli & Aly, 2020) This high performance was a result of overtraining, which was necessary to ensure matched art/room task performance across the memory-guided and explicitly instructed conditions. Second, because the attentional state selection occurred at the start of the subsequent trial, after an approximately 6.66s inter-trial interval (ITI), RTs may not be particularly sensitive measures — participants had extensive time to prepare a response during the ITI. For these reasons, neither accuracy nor RT were ideal measures for assessing the relationship between DAN-hippocampus background connectivity and the ability to retrieve memories of the stay/switch cues.

Our study is also limited in that we found a surprising null effect. Although the basal forebrain showed increased background connectivity with the hippocampus during explicitly instructed vs. memory-guided attention, as hypothesized, background connectivity strength did not predict behavior in either condition. This was unexpected because, to the extent that such connectivity enhances the processing of external signals, stronger background connectivity should predict better identification of art/room matches, perhaps in both the memory-guided and explicitly instructed conditions. One possible reason for this null effect is the difficulty of the art and room attention tasks; high hippocampus-basal forebrain background connectivity may have put individuals in a strong externally oriented state, but they may have nevertheless failed to identify art/room matches because these tasks were relatively difficult. Future research can explore the impact of basal forebrain-hippocampus background connectivity on behavioral measures of external attention in tasks in which poor performance is most likely to result from mind wandering rather than task difficulty.

One potential criticism of our approach is that the memory-guided condition may have required more task switching, or been more difficult than the explicitly instructed condition. This could complicate condition comparisons. However, four lines of evidence argue against an interpretation of our results in terms of task difficulty or generic task switching costs independent from the need to balance external and internal attention.

First, there was no difference in behavioral performance (art/room match detection) across the memory-guided and explicitly instructed conditions for valid trials, the trials that were analyzed in the current paper (t(28) = 1.32, p = 0.20, d = 0.25, 95% CI [0.058, 0.012], as reported in Günseli & Aly, 2020). Thus, extensive pre-training in detecting stay/switch cues seems to have made the memory-guided condition comparably difficult to the explicitly instructed condition, despite the greater demand on task switching in the former condition.

Second, an explanation of our findings in terms of task difficulty cannot easily account for hippocampus-DAN and hippocampus-basal forebrain connectivity moving in *opposite directions* across conditions.

Third, the connectivity flips we observed are difficult to explain in terms of generic task switching. Although some past work has implicated the cholinergic system in switching between memory systems, external stimuli, or behavioral strategies (Furey et al., 2008; Havekes et al., 2011; Aoki et al., 2015), such a “task switching” hypothesis of cholinergic function may lead to the prediction of greater hippocampal functional connectivity with the cholinergic basal forebrain in the memory-guided condition, in which more task switching might occur. However, we observed the opposite, with reduced hippocampal functional coupling with the basal forebrain in the memory-guided vs. explicitly instructed condition. Furthermore, generic task switching functions are generally more linked to the striatal cholinergic system (Furey et al., 2008; Havekes et al., 2011; Aoki et al., 2015) and not the septohippocampal system (originating in the medial septum and diagonal band of Broca in the basal forebrain), which we examined in the current study. The latter system is more implicated in prioritizing externally oriented states specifically (Sarter et al., 2003; Hasselmo 2006; Tarder-Stoll, Jayakumar, et al., 2020; Decker & Duncan, 2020) and not attentional set shifting more generally (Tait and Brown, 2008).

Fourth, we tested whether DAN univariate activity is higher in the memory-guided vs. explicitly instructed condition, which may be expected if the former condition required more top-down attentional control, a function linked to the DAN (Majerus et al., 2012; Majerus et al., 2018). However, we found no difference in univariate activity in the DAN between conditions.

Together, these lines of evidence are consistent with our proposal that our pre-training sessions minimized demands on effortful top-down control in the memory-guided condition, and that the connectivity switches we observed are more likely to be due to differential demands on external and internal attention, rather than task difficulty.

The demand to switch between attending to the stream of images and identifying stay/switch cues was indeed an important difference between our conditions — a difference that was a design feature and not a confound. The need to switch between these task components was likely important in our findings. Indeed, we propose that DAN-hippocampus background connectivity may allow for the efficient retrieval of stay/switch cue memories in the memory-guided attention task, thus freeing up attentional resources to process external visual signals and perform accurately on art/room match detection (Figure 4C).

All that said, it is possible that there were differences in task difficulty that were not detected by our behavioral measures and led to unexpected effects on connectivity patterns (although we argue this can be true of any study that finds no differences in behavioral performance). It will therefore be important to replicate our findings in future work with a different dataset. For such replications, multiple constraints must be met to allow a comparison of internal vs. external attentional states in hippocampal networks: (1) the task must include a condition that requires proportionally more internal attention, and proportionally less external attention, than a second condition; (2) behavioral performance must be balanced across conditions to avoid confounds of task difficulty; (3) the stimuli and motor demands must be identical to avoid confounds of those features; and (4) the task must involve the hippocampus. One approach may involve parametrically manipulating demands on internal attention while holding external attention demands constant, an approach we are taking in ongoing work by varying how far into the future individuals have to predict based on memory, while keeping encoding demands constant; see Tarder-Stoll, Baldassano*, & Aly*, 2023 for a similar paradigm). Such a paradigm may address limitations of the approach used here, particularly if participants are trained so that there are no differences in performance across conditions, and task switching demands are held constant.

Finally, this study was designed to compare memory-guided and explicitly instructed conditions rather than interactions between these conditions and attentional states (art or room). Future experiments can explore whether hippocampal coupling with the basal forebrain and DAN varies both by the nature of attentional cueing (from memory or external signals) and by the content of attentional states.

## Conclusion

We explored how the hippocampus may shift between external attention to the outside world and internal attention to retrieved memories. We found that the hippocampus shows stronger functional connectivity with the cholinergic basal forebrain during attention tasks that are guided by external cues rather than retrieved memories. Conversely, the hippocampus exhibited stronger functional connectivity with the dorsal attention network during memory-guided vs. externally cued attention, and this functional connectivity predicted trial-by-trial attentional behavior. Taken together, these results suggest that the basal forebrain and dorsal attention network may help the brain balance attention to the external and internal world by differentially communicating with the hippocampus — allowing us to efficiently guide behavior based on both sensory signals and memories.

## Acknowledgements

We would like to thank Dr. Serra Favila, Dr. Katherine Duncan, and the Alyssano Group for helpful feedback on this project. We also thank Dr. Eren Günseli for providing the dataset that is analyzed in the current work. This work was funded by an NSF CAREER Award (BCS-1844241) and a Zuckerman Institute Seed Grant (CU-ZI-MR-S-0001) to M.A.

